# Characterization of the Glutathione Redox State in the Golgi Apparatus

**DOI:** 10.1101/2024.12.06.627163

**Authors:** Carla Miró-Vinyals, Sarah Emmert, Gina Grammbitter, Alex Jud, Tobias Kockmann, Pablo Rivera-Fuentes

**Affiliations:** Department of Chemistry, University of Zurich, Zurich, Switzerland; Institute of Chemical Sciences and Engineering, École Polytechnique Fédéral de Lausanne, Lausanne, Switzerland; Department of Biochemistry, University of Zurich, Switzerland; Functional Genomics Center Zurich, ETH Zurich/University of Zurich, Zurich, Switzerland

**Keywords:** Golgi apparatus, roGFP, redox homeostasis, glutathione

## Abstract

Redox homeostasis is crucial for cell function, and, in eukaryotic cells, studying it in a compartmentalized way is essential due to the redox variations between different organelles. The redox state of organelles is largely determined by the redox potential of glutathione, *E*_GSH_, and the concentration of its reduced and oxidized species, *[GS]*. The Golgi apparatus is an essential component of the secretory pathway, yet little is known about the concentration or redox state of GSH in this organelle. Here, we characterized the redox state of GSH in the Golgi apparatus using a combination of microscopy and proteomics methods. Our results prove that the Golgi apparatus is a highly oxidizing organelle with a strikingly low GSH concentration (*E*_GSH_ = – 157 mV, 1–5 mM). These results fill an important gap in our knowledge of redox homeostasis in subcellular organelles. Moreover, the new Golgi-targeted GSH sensors allow us to observe dynamic changes in the GSH redox state in the organelle and pave the way for robust characterization of the Golgi redox state under various physiological and pathological conditions.

## Introduction

Redox homeostasis^1^ plays a key role in cellular signaling,^2,3^ development,^4^ and pathology.^5^ Redox imbalances are closely related to dysfunctions such as cancer,^6^ diabetes,^7^ and neurodegenerative diseases.^8^ Since the most abundant redox pair in the cell is the tripeptide glutathione (GSH) and its oxidized counterpart (GSSG),^9^ it can be assumed that GSH/GSSG is the main redox buffer in the cell. The redox state is commonly characterized by the electrochemical half-cell reduction potential (*E*_GSH_) and the total concentration of the redox pair (*[GS]*). *E*_GSH_ is typically calculated from the equilibrium concentrations of GSH and GSSG using the Nernst equation (eq.1), in which *E*°’GSH is the standard reduction potential of GSH, *R* is the gas constant (8.315 J K^-1^ mol^-1^), *T* is the temperature, and *F* is the Faraday constant (96485 C mol^-1^).

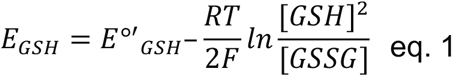

In eukaryotic cells, the redox state of GSH varies widely between organelles, indicating that redox homeostasis is compartmentalized.^10–12^ Previous studies have shown that, in HeLa cells, mitochondria are highly reducing organelles (*E*_GSH_ = –360 mV),^13^ even more so than the whole cell (*E*_GSH_ = –320 mV).^14^ In contrast, the endoplasmic reticulum (ER) is substantially more oxidizing (*E*_GSH_ = –208 mV).^15^ Moreover, several groups have quantified the GSH:GSSG ratio and the absolute concentration of GSH and GSSG or GSH alone (Figure 1).^16–21^

**Figure 1.**
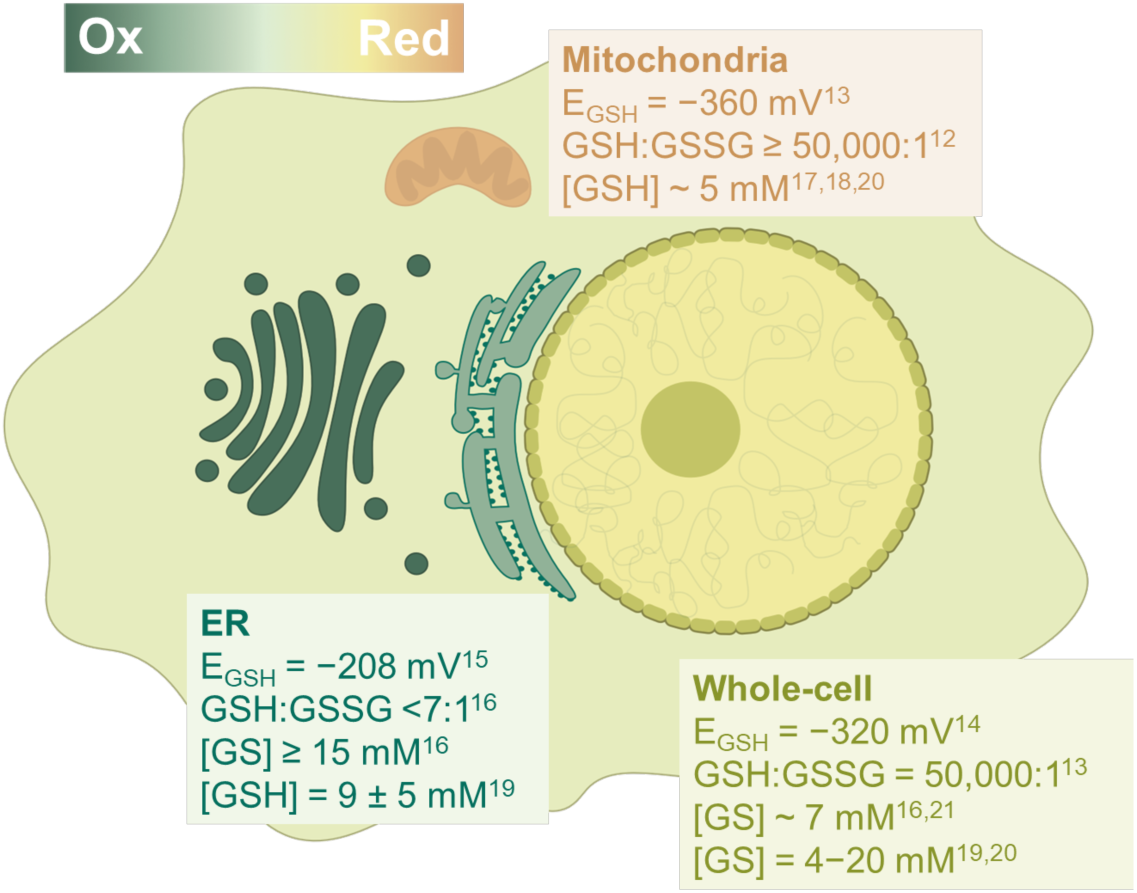
Representation of the redox state in the organelles with literature values for the *E_GSH_*, GSH:GSSG, and *[GS]*.

The Golgi apparatus (Golgi) is an organelle of the secretory pathway responsible for protein glycosylation and sorting.^22^ Despite the central role of the Golgi in the cell, the GSH redox state in this organelle remains unknown, and very little is known about thiol regulation in the Golgi in general. The only known Golgi-resident sulfhydryl oxidase or oxidoreductase in mammalian cells is quiescin sulfhydryl oxidase 1 (QSOX1).^23,24^ QSOX1 regulates the activity of sialyltransferases, most notably α-2,6- sialyltransferase (ST6Gal1), by promoting the formation of a disulfide bond once the enzyme enters the Golgi.^25^ Of note, yeast cells contain Golgi-localized glutaredoxin proteins Grx6 and Grx7,^26^ which can reduce GSSG to GSH, but there is no evidence of analogous enzymes being present in mammalian cells.

Like other organelles, redox stress in the Golgi directly impacts cellular fitness. For example, Golgi-dispersing agents induce the accumulation of lipid peroxides, a decrease in the intracellular GSH pool, and expression level changes of several ferroptosis signaling components, leading to ferroptosis-driven cell death.^27^ Moreover, Golgi-fragmentation is linked to oxidative stress and lipid peroxidation,^28,29^ and the term “glyco-redox” has been introduced to associate altered glycosylation with oxidative stress generated by hypoxia or reactive-oxygen species.^30^ All these Golgi-and redox-related alterations have been observed in disorders such as Alzheimer’s, Parkinson’s, amyotrophic lateral sclerosis, and chronic obstructive pulmonary disease,^31,32^ but further investigations are needed to establish the cause-effect relationship between the two. Therefore, understanding the basal redox state in the Golgi and developing improved tools for monitoring its GSH redox state could illuminate the stress and disease pathways activated by redox imbalance in the Golgi.

In this work, we determined the redox state and absolute concentrations of the GSH/GSSG pair in the Golgi of HeLa (cancerous) and HEK293 (non-cancerous) cells. Towards this goal, we first developed genetic and chemigenetic tools specifically tailored to study GSH in the Golgi and then applied these novel tools in fluorescence imaging and proteomics experiments to obtain robust, quantitative measurements.

## Results and Discussion

### The GSH redox potential in the Golgi is highly oxidizing

The jellyfish green fluorescent protein (GFP) has been engineered to obtain redox-sensitive GFP (roGFP) by replacing S147 and Q204 with cysteines.^33,34^ These mutations allow for the formation of a disulfide bond under oxidizing conditions that induces a change in the spectroscopic properties of the fluorophore.^29^ Hence, the excitation wavelength maximum of roGFP changes from violet (405 nm) to blue or cyan (445 or 488 nm) upon reduction. Fusing roGFP with glutaredoxin (Grx) gives roGFP kinetic selectivity for GSH/GSSG over other thiols.^34,35^ To determine the GSH/GSSG redox state in the Golgi, we developed a Golgi-targeted roGFP based on roGFP1-iE, a variant of roGFP with a dynamic range that is suitable for oxidizing environments.^15,36^ Thus, our final construct consisted of a fusion of Grx1 for GSH specificity, roGFP1-iE, for oxidizing environments, and a fragment (1-61 amino acids) of β-1,4-galactosyltransferase 1 (B4GALT1) to target the sensor to the luminal side of the Golgi membrane (Figure 2A and B). Transient expression of this construct gave a sensor that localized correctly to the Golgi (Figure 2D and Figure S1).

**Figure 2.**
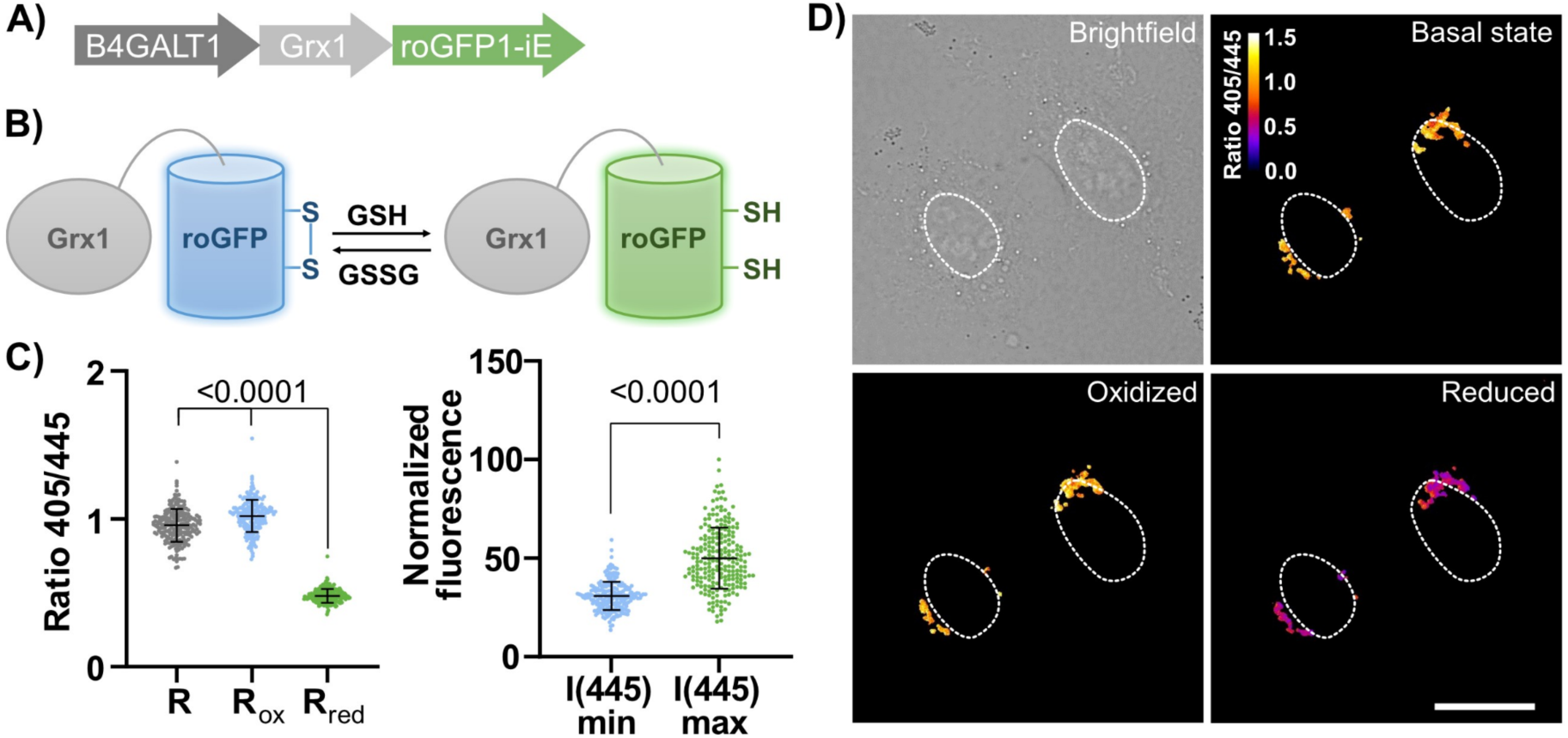
A) Gene construct for Grx1-roGFP1-iE expression in the Golgi. B) Simplified schematic equilibrium between Grx1-roGFP1-iE with GSH and GSSG. C) Data analysis of n ≥ 50, technical replicates (individual cells), average of three biological replicates. Statistical significance was assessed by unpaired, two-tailed, Mann- Whitney test. D) Representative brightfield and ratiometric fluorescent images with excitation at 405 nm and 445 nm wavelength and emission at 520/50 nm. The cells were imaged at basal state conditions, under oxidizing conditions (3 min of 1 mM H2O2 treatment), and under reducing conditions (3 min of 5 mM DTT treatment). White circles are nuclear outlines. Scale bar = 20 µm.

By assuming an equilibrium between roGFP1-iE and GSH, we calculated the *E*_GSH_ by using the modified Nernst equation (eq. 2), in which *E°’_roGFP1-iE_* is obtained using eq. 3 and considering a pH value for the Golgi of 6.2 (*E°’_roGFP1-iE, pH 6.2_* = –190 mV). *OxD*_roGFP_ (eq. 4) is the degree of oxidation of the sensor, and it can be calculated by measuring the fluorescence emission intensity at 520 nm with excitation at 405 nm (oxidized) and 445 nm (reduced).^13,15,37^ Fluorescence ratios of excitation at 405/445 were measured for the basal state (*R*), under reduced conditions (*R_red_*, upon treatment with dithiothreitol (DTT)), and under an oxidized environment (*R_ox_*, H_2_O_2_ treatment) (Figure 2C). Finally, by measuring the fully reduced and fully oxidized fluorescence intensities at 445 nm, (*I(445)_max_* and *I(445)_min_*, respectively) one obtains *OxD_roGFP_*.

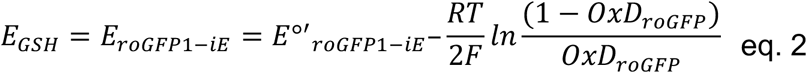

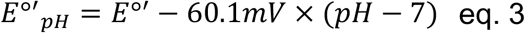

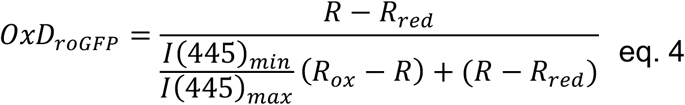

By measuring cells at 520/50 nm fluorescence emission with 405 and 445 nm excitation wavelengths, we obtained *E*_GSH_ = –157 ±3 mV for the Golgi. We performed the same experiment for HEK293 cells (Figure S1) obtaining a value of *E*_GSH_ = –147 ±4 mV. These values indicate that the Golgi in either cell type has a more oxidizing environment than the ER in HeLa cells (*E*_GSH_(ER) = –208 ±4 mV) using the same fluorescent sensor.^16^

### The ratio of GSH to GSSG in the Golgi is much lower than in the cytosol

Although the redox potential of the GSH pair depends on the GSH:GSSG ratio, the latter cannot be calculated directly from the *E*_GSH_ value without knowledge of the absolute concentrations. To obtain the GSH:GSSG ratio, we employed a mutant of the yeast glutaredoxin Grx1 in which one of the two cysteine residues was mutated to serine (sCGrx1p).^16^ Having only one reactive cysteine, sCGrx1p can equilibrate rapidly and reliably with the GSH:GSSG pool in the cell (Figure 3B) with an equilibrium constant *K*ox of 234 ±6 at pH 6.2.^38^ Quantification of the GSH:GSSG ratio (*RGS*) can be, thus, obtained using eq. 7, provided that the glutathionylated peptide of sCGrx1p can be quantified by, e.g., liquid chromatography coupled to tandem mass spectrometry (LC-MS/MS).

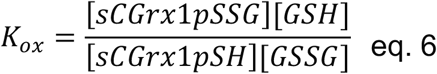

**Figure 3.**
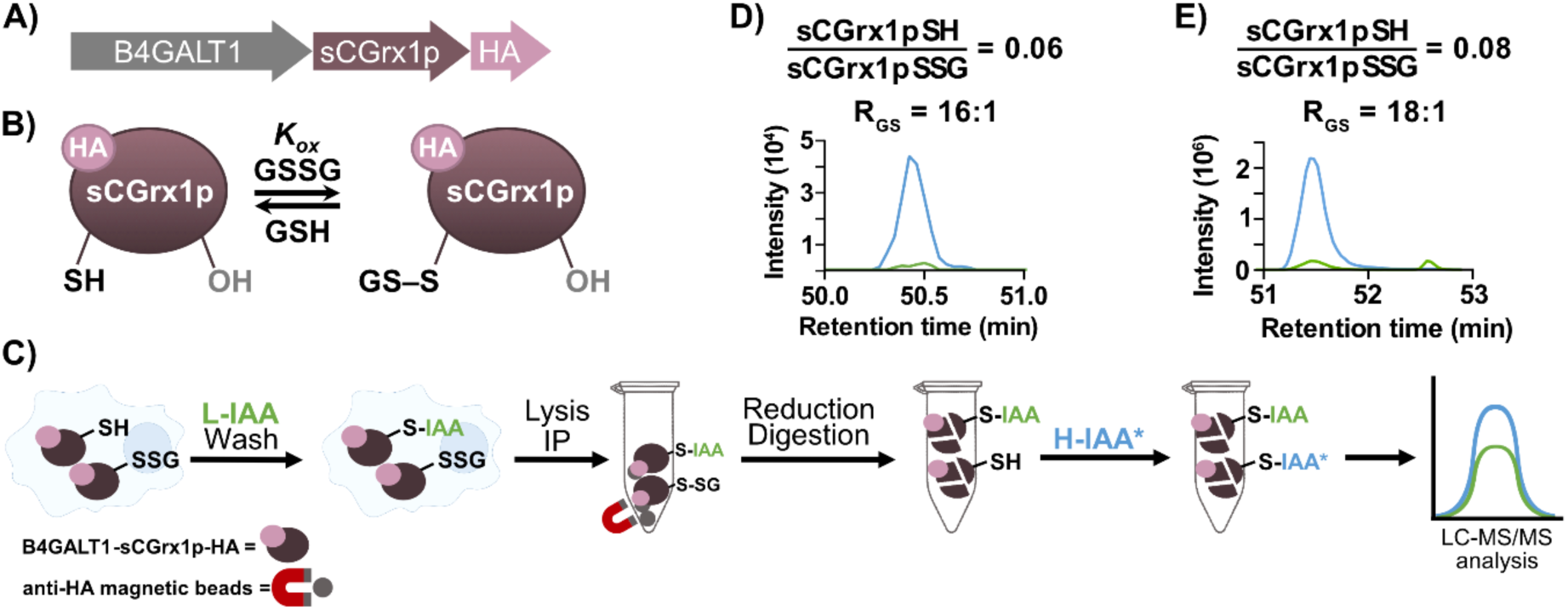
A) Gene construct for sCGrx1p-HA expression in the Golgi. **B)** Schematic equilibrium between sCGrx1p with GSH and GSSG. **C)** Protein derivatization workflow for targeted proteomics analysis of degree of glutathionylation of sGCrx1p. **D)** and **E)** Extracted-ion chromatogram (XIC) trace in MS2 of the heavy and light Cys-containing peptide (TYCPYSHAALNTLFEK) of fragment ion y14++ (825.9009++) and quantification to *RGS*. **D)** HeLa cells lysate, **E)** HEK293 cells lysate. Green color indicates the reduced sensor protein/light derivate. Blue indicates the oxidized sensor protein/heavy derivative. Reactive Cys of sensor protein is underlined in the peptide sequence above.

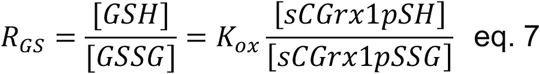

Using recombinant protein in vitro, we demonstrated that sCGrx1p is quantitatively glutathionylated by GSSG (Figure S2). Additionally, we confirmed that glutathionylation can be reversed using tris(2-carboxyethyl)phosphine (TCEP), allowing the reduced protein to be tagged at the relevant cysteine with an alkylating agent such as *N*-ethylmaleimide (NEM) or iodoacetamide (IAA) (Figure S2). This reactivity opens the opportunity for quantitative LC-MS/MS determination of the reduced (sCGrx1pSH) and glutathionylated (sCGrx1pSSG) states of the protein by alkylation with isotopically labeled reagents.

We developed a Golgi-targeted (B4GALT1) sCGrx1p probe equipped with a human influenza hemagglutinin (HA) peptide tag for immunoprecipitation (IP). The probe was encoded in a plasmid for transient transfection (Figure 3A and Figure S3). The correct expression and localization of the B4GALT1-sCGrx1p-HA probe were assessed by immunofluorescence against the Golgi-resident protein GM-130 (Figure S3). Finally, the alkylating agent of choice was IAA due to better peptide MS/MS selectivity (Figure S3).

To obtain *RGS* (eq. 7), we quantified the degree of cysteine glutathionylation of sCGrx1p (*[sCGrx1pSH]* and *[sCGrx1pSSG]*) in the Golgi by a targeted proteomics approach using light and heavy iodoacetamide labeling (L- and H-IAA, respectively). The workflow (Figure 3C) consisted of alkylating the free-thiol fraction of sCGrx1p (sCGrx1pSH) in the Golgi of live cells with L-IAA (ICH2CONH2) followed by extensive washing, and subsequent cell lysis, and protein enrichment by IP with anti-HA beads. Reduction of the glutathionylated protein (sCGrx1pSSG) with TCEP, followed by on-bead digestion and final labeling of the newly formed cysteine with H-IAA (I^13^CD2^13^CONH2) yielded the sample for LC-MS/MS analysis. The unmodified form was also monitored, but the corresponding fragment ion signals were always below the limit of detection.

The pipeline resulted in a ratio of [sCGrx1pSH]/[sCGrx1pSSG] of 0.06, which corresponded to an *RGS* of 16:1 (Figure 3D) in HeLa cells. In HEK293 cells, the obtained [sCGrx1pSH]/[sCGrx1pSSG] ratio was 0.08, which corresponded to an *RGS* of 18:1 (Figure 3E).

The fact that the redox potential indicates the Golgi as a more oxidizing organelle than the ER, whereas the GSH:GSSG ratio suggests that the Golgi has fewer GSSG equivalents than the ER (*RGS* < 7)^16^ might seem contradictory at first. However, this observation would be consistent with a lower concentration of GSH in the Golgi compared to the ER, since the redox potential of GSH depends not only on the GSH:GSSG, but also on the concentration of GSH (eq. 1).

Finally, by using the above obtained *E*_GSH_ (also represented as *[GSH]^2^*:*[GSH]*, with a value of 81 mM for HeLa cells and a value of 37 mM for HEK293 cells) and the *RGS* values obtained in this section, we calculated the total concentration of GSH and GSSG in the Golgi (*[GS]*) by using eq. 8 and 9.

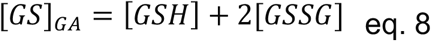

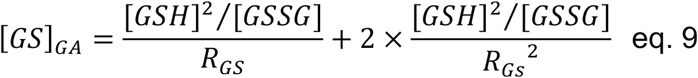

The *[GS]* in the Golgi of HeLa cells was determined as 5 mM, whereas in HEK293 cells a value of 2 mM was obtained. The concentration of GSH and GSSG in the Golgi resulted in a much lower concentration than for the ER, when calculated with the same sCGrx1p and roGFP1-iE sensors (> 15 mM).^16^

### The concentration of GSH in the Golgi is particularly low

To confirm that the Golgi has an unusually low concentration of GSH and GSSG, we employed an alternative strategy. We recently reported the development of a chemigenetic, quantitative, fluorescent sensor for GSH, which we called TRaQ-G.^20^ The sensor consists of a fusion protein of HaloTag (HT) – a self-labeling protein – and mGold – a yellow fluorescent protein with high brightness. HT conjugates the TRaQ- G ligand, a GSH-reactive silicon rhodamine (SiR) dye. This method takes advantage of the fluorogenicity of SiR, which only becomes fluorescent – and GSH-reactive – when bound to HT (Figure 4B), hence solely sensing GSH in the selected location in which the protein is expressed. Moreover, the GSH-insensitive fluorescent internal standard mGold makes the sensor suitable for absolute GSH-concentration determination via ratiometric measurements. We developed a Golgi-targeted TRaQ- G sensor using the previously used B4GALT1 fragment (Figure 4A and Figure S4). Furthermore, we tested the functionality of the sensor by subjecting cells to conditions that changed the endogenous concentration of GSH (Figure 4E). The ethyl ester of GSH (EtGSH), which is membrane permeant, was used to induce an increase in the concentration of GSH, giving an increase in the mGold/TRaQ-G ratio, as expected. To lower the GSH concentration, we used buthionine sulfoximine (BSO), an inhibitor of γ- glutamylcysteine synthetase, a key enzyme in the biosynthetic pathway of GSH (Figure 4E). In this case, the mGold/TRaQ-G ratio decreased, as expected. With this experiment, we prove that the Golgi-targeted sensor is responding to changes in the GSH pool of the organelle. Of note, since the induced increase and decrease of GSH originates in the cytosol – BSO inhibits a cytosolic enzyme, while the GSH ester is cleaved by esterases in the cytosol – the experiment also proves that GSH, and possibly GSSG, are transported across the Golgi membrane since the Golgi GSH pool responds to extreme changes originating in the cytosol.

**Figure 4.**
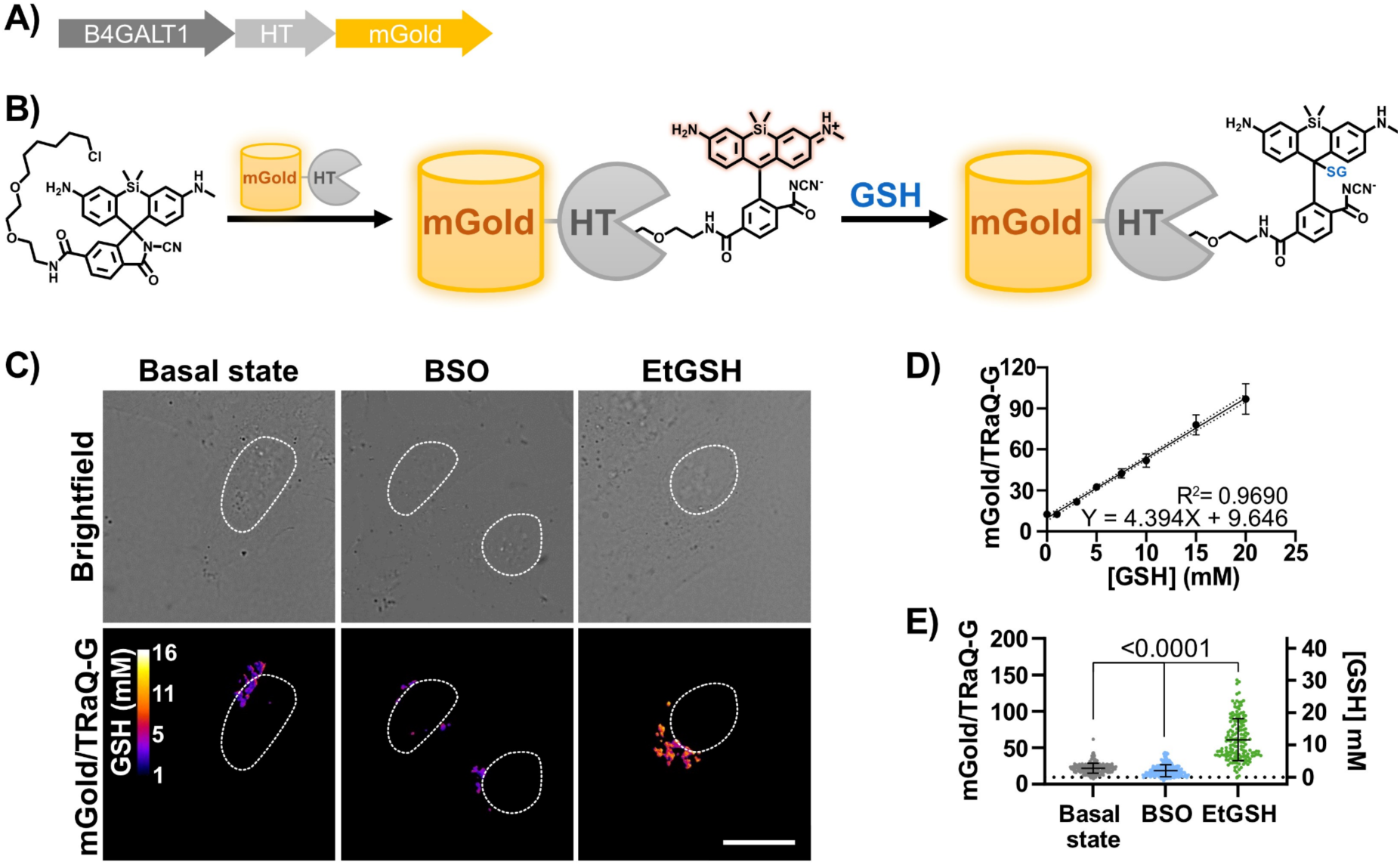
A) Gene construct for HT-mGold expression in the Golgi. **B)** Schematic representation of the working principle of the TraQ-G-mGold construct. **C)** Brightfield and ratiometric fluorescent images of HeLa cells transiently transfected with B4GALT1-HT-mGold and incubated with 100 nM TRaQ-G ligand for 1 h at 37 °C. Excitation at 515 nm and emission at 542/27 nm for mGold, and excitation at 561 nm and emission at 642 long-pass filter for TRaQ-G. The cells were imaged at basal state conditions, under BSO treatment (1 mM, 3–4 h incubation) and under EtGSH incubation (10 mM, 3–4 h incubation). White circles are nuclear outlines. Scale bar = 20 µm. **D)** Data analysis of the *in vitro* calibration curve with purified HT-mGold protein conjugated to TRaQ-G and incubated with increasing amounts of GSH. Imaged with confocal microscopy with identical optical configuration as for living cells. **E)** Data analysis of n ≥ 50, technical replicates (individual cells), average of three biological replicates. Statistical significance was assessed by unpaired, two-tailed, Mann-Whitney test.

To quantify the concentration of GSH in the Golgi, we expressed and purified an HT- mGold-6His construct and conjugated it to the TRaQ-G ligand. We used this sensor to generate a calibration curve relating mGold/TRaQ-G ratios to GSH concentrations under the same imaging conditions of live-cell experiments (Figure 4D).

Our results indicate that the GSH concentration in the Golgi is rather low, at a range of 2.7 ±1.5 mM values in HeLa cells (Figure 4E). This figure contrasts with substantially higher concentrations in the ER (9 ±5 mM), nuclei (14 ±7 mM), or mitochondria (4.8 ±0.7 mM), measured with the same sensor.^20^ The concentration of GSH in the Golgi of HEK293 cells was even lower, at 1.1 ±1.5 mM (Figure S4).

Despite the contrast to the higher concentrations of GSH in the ER, the results from TRaQ-G agree with the results obtained with sCGrx1p, which also show a low mM concentration of total GSH and GSSG in the Golgi.

## Conclusions

We developed several tools to characterize the GSH redox state in the Golgi, two of which are based on a fluorescence readout. We found that the Golgi’s GSH redox potential is –157 ±3 mV for HeLa cells and –147 ±4 mV for HEK293 cells with roGFP sensors. These measurements reveal that the Golgi is substantially more oxidized than other organelles, such as the nucleus, mitochondria, or even the ER. Moreover, we quantified the GSH:GSSG ratio and the absolute concentration of GSH and GSSG in the Golgi with sCGrx1p, being 16:1 and 5 mM in HeLa cells and 18:1 and 2 mM in HEK293 cells, indicating a highly oxidized *RGS*, and a *[GS]* lower than in any other quantified organelle thus far (including the ER).^9,20^ Independently, TRaQ-G allowed for the determination of an absolute GSH concentration in the Golgi of 2.7 ±1.5 mM for HeLa cells and 1.1 ±1.5 mM for HEK293 cells, which are in the same low mM range as obtained with sCGrx1p. We are confident that the values we obtained are reliable and robust by utilizing three distinct techniques based on different GSH detection principles (roGFP, sCGrx1p, and TRaQ-G). To the best of our knowledge, these results provide the first reference values for the redox state and concentration of GSH/GSSG in the Golgi of cancerous and non-cancerous cells (Table 1).

**Table 1.** Calculated GSH redox state values in the Golgi apparatus of HeLa and HEK293 cells.

|  | HeLa cells | HEK293 cells |
| --- | --- | --- |
| $E_{GSH}$ | −157 mV | −147 mV |
| $[GSH]^2:[GSSG]$ | 81 mM | 37 mM |
| $R_{GS}$ | 16:1 | 18:1 |
| $[GS]_{sCGrx1p}$ | 5.8 mM | 2.1 mM |
| $[GSH]_{TRaQ-G}$ | $2.7 \pm 1.5$ mM | $1.1 \pm 1.5$ mM |

Open questions beyond the scope of this study include why a more oxidizing redox potential is needed in the Golgi than in the ER, or how GSH and GSSG are transported into or from the Golgi – and whether synthesis, degradation, oxidation, or reduction of GSH/GSSG occur in the Golgi. We envision that our newly tailored tools will be instrumental in testing these hypotheses by monitoring intra-organelle fluctuations of GSH/GSSG concentrations in the Golgi with spatiotemporal resolution. Therefore, this research opens new avenues to study redox homeostasis in an important, yet largely underexplored organelle.

## Methods

Unless stated otherwise, all reagents and solvents were purchased from commercial sources and used as received.

### Mammalian cell culture

HeLa or HEK293 cells were grown in Dulbecco’s Modified Eagle Medium (DMEM, low glucose 1 g L^−1^, Qiagen #31885-023) supplemented with 10% fetal bovine serum (FBS, Gibco #A5256801) 100 U mL^−1^ penicillin, 100 µg mL^−1^ streptomycin and 0.25 mg mL^−1^ Fungizone at 37 °C in 5% CO2 environment. The cells were passed after 90% confluence was reached and they were split a maximum of 15 times.

### Transient plasmid DNA transfection

For imaging, 15’000 – 30’000 cells were seeded per well of an 8-well Ibidi chambered cover glass 48 to 72 h prior to imaging. Cells were transfected with plasmid DNA using jetPRIME^®^ according to the recommended protocol of the supplier 24 to 48 h prior to imaging. If specified, the cells were incubated with the respective small-molecule probes in growth medium for the indicated time. Before imaging, the growth medium was removed, the cells were washed 2x with PBS and imaged in FluoroBrite DMEM (Qiagen #A1896702).

### Cell fixation and immunofluorescence

Cells were grown for 24 to 48 h on an Ibidi 8-well-plate, fixed with 4% paraformaldehyde for 20 min at 20°C, washed twice with phosphate-buffered saline (PBS), and permeabilized with 0.1% Triton X-100 for 15 min at 20°C. After three washing steps, the cells were blocked with 2% bovine serum albumin (BSA) in PBS for one hour, with rotation, at 20 °C. The cells were incubated in 0.1% BSA in PBS with mouse anti-HA (ThermoFischer, 26183, 2 µg mL^−1^ dilution) and/or rabbit anti-GM130 (ThermoFischer, PA5-95727, 1:100 dilution). After 16 h incubation at 4°C, three washing steps with PBS were performed for 20 min each. Then, the secondary antibodies goat anti-mouse-AlexaFluor488 (ThermoFischer, A32723, 10 µg mL^−1^ dilution) and/or donkey anti-rabbit-AlexaFluor680 were added to the cells (Abcam, ab186692, 16 µg mL^−1^ dilution). The incubation proceeded for 1 h at 20 °C. After several washes with PBS, the cells were imaged with a Nikon W1 spinning disc microscope. See Table S1 for details of antibodies. No bleed-through between fluorescence channels and no cross-labeling between the antibodies was observed.

### Confocal microscopy and image analysis

Imaging was performed with a Nikon W1 spinning disk microscope equipped with a dual-camera system (CMOS, Photometrix). Brightfield imaging was performed with a white LED. Available laser lines for fluorescence imaging were 405 nm, 445 nm, 488 nm, 514 nm, 561 nm, and 638 nm. Appropriate emission filters were configured within the light path as described in Table S2. Images were collected using a CFI Plan Apochromat Lambda D oil immersion objective (60x, NA = 1.4), a CFI Plan Apo VC water immersion objective (60x magnification, NA = 1.2), or a CFI SR HP Apo TIRF oil immersion objective (100x magnification, NA = 1.49). The microscope was operated using the NIS elements software. Multiple channel images were acquired by sequential imaging.

General image processing was conducted with the Fiji – ImageJ software, and image processing for data analysis was conducted with the CellProfiler software^39,40^ (Data analysis pipelines can be found in DOI: 10.5281/zenodo.14288576). Generally, the background from images was corrected using a minimum pixel intensity in a 600-pixel block size, and later a Gaussian filter smoothing method. Golgi objects were selected by first identifying cell objects with a Robust Background thresholding method. Then, after enhancing the Golgi spots in the image and masking them, applying a second object-identifying step the Golgi is selected and grouped together per cell with the selected cell objects. This is done for the typical two channels for roGFP and TRaQ- G sensors, and finally, an image and result data set made from the division of the two channels, and the Golgi objects are generated. All data were plotted using GraphPad – Prism 9.3.1 and statistical significance was assessed by unpaired, two-tailed, Mann-Whitney test.

### Gibson cloning

Primers for amplification (minimum 15 overlapping base pairs) were generated using SnapGene® and modified manually to minimize hairpins, homodimers, and repeating motifs. Primers were supplied by Mycrosynth AG (Switzerland). Respective precursor plasmids and primers are reported in Table S3 and Table S4. Reagents and general procedures from the Gibson Assembly Cloning Kit from New England Biolabs (NEB) were used. Vector and insert fragments were linearized by PCR using Phusion High-Fidelity PCR Master Mix with HF (NEB #M0531S), and template DNA was digested with DpnI (NEB #R0176S). The fragments were analyzed by gel electrophoresis. PCR fragments were purified with the QIAquick PCR Purification Kit (Qiagen, #28104). Gel extraction was pursued if the PCR purity was not sufficient with the Gel Extraction Kit (Qiagen #28706X4). Backbone and insert fragments were ligated using Gibson Assembly Master Mix (NEB #E2611S) and the product was used to transform NEB 5α competent E. coli cells (NEB #C2987H) by heat shock following the provided standard procedure of the vendor and streaked onto LB agar plates containing the corresponding antibiotic. After incubation at 37°C for 24 h, single colonies were selected and grown in LB liquid medium containing the corresponding antibiotic at 37°C for 16 h. Plasmid DNA was extracted and purified using a QIAprep spin mini or midi-prep kit (Qiagen #27106 or #12943) according to the manufacturer’s instructions and sequenced by Mycrosynth AG (Switzerland). When needed, plasmids were amplified by incubation of LB cultures with appropriate antibiotics overnight at 37 °C, DNA isolated, and sequenced as mentioned above.

### Protein expression

The protein expression plasmid was transformed into BL21-DE3 cells (NEB # C2527H). The cells were streaked onto LB agar plates containing the corresponding antibiotic. After incubation at 37°C for 24 h, single colonies from each transformation were used to inoculate 80 mL pre-cultures of Lysogeny broth (LB) medium with suitable antibiotics. The pre-cultures were shaken at 37°C, at 180 rpm for 16 h. 20 mL of the pre-cultures were used to inoculate a 1 L solution of LB medium (with the appropriate antibiotic) to yield protein-expression cultures for each replicate. The cultures were shaken at 37 °C and 180 rpm until optical density OD600 reached 0.6 – 0.8 (approx. 2h). The temperature was decreased to 18 °C and protein expression was induced by the addition of IPTG (1 mM). The cells were shaken at 18 °C for 18 h. An SDS-PAGE (4–20% Mini-PROTEAN® TGX™, Bio-Rad #4568093) was performed to check for protein expression before IPTG induction and after 18 h of induction. The cells were harvested by centrifugation (5000 xg for 15 min at 4 °C) and the pellets were collected for protein purification.

### Protein purification with Ni-NTA resin

The pellet was resuspended with wash buffer (PBS, pH 7.2) and 10% glycerol, turbonuclease (AG Scientific T-1222), and protease inhibitor (cOmplete™ Mini EDTA-free Protease Inhibitor Cocktail Roche #73953000) were added. Cells were lysed by sonication (2.5 min, 10 s, and 10 s pulse, 70% power). After centrifugation (20000 xg, 30 min, 4 °C), the supernatant was filtered through a 0.45 µM filter and incubated in batch mode with His-Pur Ni-NTA resin (ThermoFisher) + 20 mM imidazole (3 h, rotating, at 4 °C).

The resin with the protein-bound was loaded in a column and subsequent washing and eluting steps were performed:

- Wash with 20 mM imidazole, PBS, 7.5 pH - 4x25 mL
- Wash with 50 mM imidazole, PBS, 7.5 pH - 4x12.5 mL
- Wash with 100 mM imidazole, PBS, 7.5 pH - 2x12.5 mL
- Elute with 300 mM imidazole, PBS, 7.5 pH - 5x10 mL
- Elute with 500 mM imidazole, PBS, 7.5 pH - 1x15 mL

An SDS-PAGE (4–20% Mini-PROTEAN® TGX™, Bio-Rad #4568093) was run according to the supplier’s protocol to check for the purity of the fractions. The fractions that showed pure and high concentration protein were pooled together and set to dialysis against PBS for 16 h at 4 °C. The concentration was measured by measuring the absorbance at 500 nm and, if necessary, the samples were concentrated.

### UPLC-ESI-MS experiments of sCGrx1p, alkylation

Purified sCGrx1p (94 µM, 50 µL) was incubated with NEM or IAA (20x concentrated, 2 mM from the 50 mM stock, 5% DMSO) at 4 °C, for 16 h. For MS preparation, the sample was diluted to 30 µM and 1% formic acid (FA) was added. 10 µL of the sample was filtered through a C18 ZipTip (Merck Milipore) according to the supplier’s protocol (elution with 20 µL ACN/H2O/FA, 75:25:0.1).

About 1 pmol to 5 pmol of protein samples were injected into the Xevo G2-XS QTOF mass spectrometer (Waters). The samples were separated using a 10% to 90% gradient (buffer A 0.1% FA in ddH2O, buffer B 0.1% FA in ACN) over 6 min to 15 min on an Acquity UPLC Protein BEH C4 column (Waters). Eluted proteins were ionized by electron-spray ionization (ESI) in positive mode and analyzed using the following settings: source temperature 100 °C, desolvation temperature 400 °C, capillary voltage 3 kV, cone voltage 80 kV, and source offset 50 kV. The raw spectra were analyzed with Mnova MSChrom.

### UPLC-ESI-MS experiments of sCGrx1p, GSH/GSSG titration

Purified sCGrx1p (94 µM, 50 µL) was incubated with the indicated GSH/GSSG concentrations for 10 min at 20 °C. The reaction was capped by incubation with NEM (50 mM) for 16 h at 4 °C. For the TCEP-reduced proteins, 0.5 mM TCEP was added to the protein, and removal was performed with Zeba™ Spin Desalting Columns (7K MWCO, 0.5 mL, ThermoFisher #89882) and Amicon® Ultra Centrifugal Filter (3 kDa MWCO, Millipore #UFC500308).

For MS preparation the sample was diluted to 30 µM and 1% formic acid (FA) was added. 10 µL of the sample was filtered through a C18 ZipTip (Merck Milipore #ZTC18S096) according to the supplier’s protocol (elution with 20 µL ACN/H2O/FA, 75:25:0.1). Same LC-ESI-MS settings as above were used.

### sCGrx1p western blot

Handcasted gels (Stacking gel: 4 % Acryl- /Bisacrylamid 29:1, Separation gel: 16 % Acryl-/Bisacrylamid 29:1, 0.6 M Urea) were run as duplicates (Coomassie staining, Western blot) in Tris-Tricine buffer at 40 V for 20 min, followed by 200 V for ca. 1 h. For the Coomassie staining, the gels were incubated for around 20 min in InstantBlue® (Abcam), followed by destaining in water. For western blotting, the proteins were transferred with the Trans-Blot® TurboTM Transfer System (Bio-Rad) to PVDF membranes (Mini Transfer Pack, 0.2 µm PVDF, #1704156) using the “Mixed MW” protocol. Membranes were blocked for 2 h at 18 °C with a solution of 5% skim milk in PBS containing 0.1% Tween 20 (PBS-T). After a subsequent washing step with PBS-T, the membranes were incubated with the primary antibodies (Mouse-anti-GAPDH conjugated with Dylight 488; 0.5 μg/ml, Rabbit-anti-HA; 1 μg/ml) diluted in 1% skim milk in PBS-T at 4 °C overnight. After three 10 min washes with PBS-T, the membranes were incubated with the secondary antibody (Donkey-anti-Rabbit conjugated with AlexaFluor 680; 0.2 μg/ml) diluted in 1% skim milk in PBS-T at 18 °C for 1 h. Membranes were washed four times for 5 min with PBS-T, followed by two times 5 min with PBS. Imaging for Western blots and Coomassie stained gels was carried out using the ChemiDoc MP System (Bio-Rad) using the suitable device settings.

### Sample preparation for sCGrx1p targeted proteomics analysis

*Cell growth and treatment.* Cells were grown as indicated before and seeded to a 100 mm dish at 3– 5M cells per dish until confluency was reached. Cells were transfected with B4GALT1-sCGrx1p-HA plasmid DNA using jetPRIME^®^ according to the recommended protocol of the supplier 24 h before treatment. Cells were then washed 2x with ice-cold HBSS, 0.1 mM iodoacetamide (IAA) and incubated in the same buffer for 20 min on ice in order to alkylate the reduced sCGrx1p protein. Immediately after, the cell monolayer was washed three times with ice-cold HBSS to remove excess IAA.

*Cell lysing.* Cells were then trypsinized (with warm 0.5% trypsin with EDTA at 37 °C for 2 min) and resuspended in full growth medium. Cells were counted with the EVE™

Automatic Cell Counter. The cells were pelleted (4 °C, 450 x g, 3 min), and washed twice with ice-cold PBS. Cells were immediately lysed by adding CytoBuster (Merck Millipore #71009-M, 150 µL/million cells) and a protease inhibitor cocktail for mammalian cells (Sigma #P8340, 1.5 µL/million cells), and let stand for 5 min at 20 °C. The suspension was transferred to an appropriate tube and centrifuged (4 °C, 16,000 xg, 10 min). The cleared lysate was used right away for IP or stored at –80 °C. Protein concentration was determined by using the bicinchoninic acid (BCA) assay following the supplier’s protocol (Novagen #712853).

*Immunoprecipitation.* The B4GALT1-sCGrx1p-HA protein was immunoprecipitated with anti-HA magnetic beads (Pierce, #88837) according to the supplier’s protocol. The sample was incubated 16 h at 4 °C for optimal binding to the beads. For proteomics, the sample bound to the beads was subjected to further modifications directly.

*On-beads digestion, and further derivatization for proteomics.* The protein was directly digested on the beads after washing twice with digestion buffer (10 mM Tris/2 mM CaCl2, pH 8.2), and the wash was discarded. Then, digestion buffer and 10% trypsin (100 ng/µl in 10 mM HCl) were added to the beads and a microwave-assisted digestion was performed (60° C, 30 min). The supernatant was collected, and the resulting peptides were extracted from beads with digestion buffer. Reduction of Cysteins was conducted by adding ∼ 2% of TCEP (100 mM stock) and incubation for 30 min at 30 °C. Afterward reduced cysteines were alkylated by addition of heavy-IAA to a final concentration of 10 mM (Merck, CAS #1619234-07-9) and incubation for 30 min, at 20 °C, in the dark.

### sCGrx1p targeted proteomics analysis

LC-MS/MS analysis was performed on an Orbitrap Fusion Lumos (Thermo Scientific) equipped with a Digital PicoView source (New Objective) and coupled to an M-Class UPLC (Waters). The solvent composition of the two channels was 0.1% formic acid in MilliQ water for channel A and 0.1% formic acid in 99.9% acetonitrile for channel B. Column temperature was 50°C. For each sample 1:10 dilution was prepared and 1 μL of the diluted peptide stock was loaded on a commercial ACQUITY UPLC M-Class Symmetry C18 Trap Column (100 Å, 5 µm, 180 µm x 20 mm, Waters) connected to a ACQUITY UPLC M-Class HSS T3 Column (100 Å, 1.8 µm, 75 µm X 250 mm, Waters).

The peptides were eluted at a flow rate of 300 nL min^−1^. After a 3 min initial hold at 5% B, a gradient from 5 to 22% B in 40 min and 22 to 32% B in additional 5 min was applied. The column was cleaned after the run by increasing to 95% B and holding 95% B for 10 min prior to re-establishing the loading condition.

The mass spectrometer was operated in targeted mode (Parallel Reaction Monitoring –PRM). Targeted MS2 scans covered the following masses (monoisotopic peptide ions): 724.8852 (DLIAENEIFVASK++), 957.9564 (TYC[+57.021464]PYSHAALNTLFEK++, light), 959.966 (TYC[+57.021464]PYSHAALNTLFEK++, heavy), 638.9733 (TYC[+57.021464]PYSHAALNTLFEK+++, light), 640.3131 (TYC[+57.021464]PYSHAALNTLFEK+++, heavy), 929.4456 (TYCPYSHAALNTLFEK++), 619.9662 (TYCPYSHAALNTLFEK+++), 586.839 (VLVLQLNDMK++), 874.4341 (EGADIQAALYEINGQR++), 559.8139 (TVPNIYINGK), and 683.8391 (HIGGNDDLQELR++). Each target ion was isolated in the Quadrupole using an isolation window of 0.8 Da centered on the target value. Targeted MS2 scans were acquired in the Orbitrap mass analyzer at 60’000 resolution, with a normalized AGC target value of 100% (equal to 50’000 charges), a maximum injection time of 118 ms and a fixed normalized collision energy (NCE) of 30% applying HCD activation. Each instrument cycle was completed by a full MS scan monitoring 350 to 1500 m/z at a resolution of 120’000 using the Orbitrap mass analyzer.

*Raw LC-MS data handling.* The raw LC-MS data were handled using the B-Fabric information management system^41^ and all relevant data have been deposited to PRIDE (identifier to be added) and Zenodo (DOI: 10.5281/zenodo.14288576).

*Targeted quantification.* PRM data analysis was carried out on Skyline by manual curation. In short, the light- and heavy-IAA Cys-containing labeled peptides (TYCPYSHAALNTLFEK++) XIC areas were obtained by analyzing the y14++ fragment (825.9009++) see Figure S3D for more details.

### TRaQ-G calibration curve

The TRaQ-G sensor was assembled in vitro using the purified HT-mGold-6His fusion protein (220 µM in PBS, pH 7.2) and incubating with 2x TRaQ-G ligand (440 µM, stock solution in DMSO) for 1 h at 20 °C. The adduct was used in a final concentration of ∼10 µM by diluting it in phosphate buffer (0.5M Na2HPO4, NaH2PO4, pH 6.6). In a 96- well plate (glass-bottom), the adduct was treated with the appropriate amount of GSH (freshly made solution in phosphate buffer). The mGold/TRaQ-G ratio was measured by fluorescence microscopy. The ratiometric measurement was carried out with the same instrument with the same settings as for the live-cell experiments. The measurement was performed with three technical replicates and the blank averaged over all concentrations. Three fields of view (FOVs) were measured per well. The image analysis was performed with Fiji – ImageJ. The background was determined from the blank measurements calculating the mean of all the pixels in the FOV per channel excluding the edges. The background was subtracted for the sample measurements and each channel. The mGold channel was divided by the TRaQ-G channel in a pixel-by-pixel manner and the mean of all pixels in the FOV excluding the edges was calculated. Every FOV gave one data point.

### GSH measurement in living HeLa cells

HeLa cells were plated and transfected with the respective HT-mGold plasmid. After 24 h, the cells were incubated with 100 nM TRaQ-G ligand for 1 h. The cells were treated with 10 mM EtGSH for 3–4 h, or 1 mM BSO for 3–4 h in FluoroBrite DMEM. GSH concentration was measured by fluorescence microscopy using the calibration curve. The experiment was carried out in three biological replicates on different days with cells from different passages. The data sets were combined for analysis. In total, 90–200 cells were analyzed per condition. The calibration curve was used to interpolate/extrapolate means as well as upper and lower bounds expressed in GSH concentration for single cells.

## Author contributions

All experiments were carried out by C. M.-V. S. E. synthesized the TRaQ-G ligand probe. G. G. assisted with the intact MS *in vitro* experiments and data analysis. A. J. performed the sCGrx1p sensor optimization. T. K. performed LC-MS/MS experiments. C. M.-V. and T. K. analyzed UPLC-ESI-MS/MS data. P. R.-F. supervised the project and raised funds. C. M.-V. and P. R.-F. conceived the project, analyzed data, and wrote the manuscript with input from all authors. All authors have read and agreed to the published version of the manuscript.

## Funding

This work was funded by the Swiss National Science Foundation (grant no. PCEGP2_186862 to P. R.-F.).

## Declaration of competing interest

None declared.

## Notes

The authors declare no competing financial interest. All the raw data from this paper are available on Zenodo with DOI: 10.5281/zenodo.14288576, proteomics data is available at PRIDE (identifier to be added prior to publication).

## Acknowledgements

We thank Gianluca Quargnali (University of Zurich) for cloning the B4GALT1-HT- mGold plasmid and staff at the Functional Genomics Center Zurich for assistance with sample preparation and acquisition for proteomics studies.

## Notes

https://doi.org/10.5281/zenodo.14288576

### Competing Interest Statement

The authors have declared no competing interest.

