## Supplementary Information for "Characterization of the Glutathione Redox State in the Golgi Apparatus"

For manuscript:

1. Supplementary table
2. Supplementary figures

### 1. Supplementary tables

**Table S1. List of antibodies.**

| Name and host | Conjugated | Supplier | Supplier number |
|---|---|---|---|
| Mouse anti-HA | No | ThermoFisher | 26183 |
| Rabbit anti-GM130 | No | ThermoFisher | PA5-95727 |
| Goat anti-mouse | Alexa 488 | ThermoFisher | A32723 |
| Donkey anti-rabbit | Alexa 680 | abcam | ab186692 |
| Mouse-anti-GAPDH | Dylight 488 | ThermoFisher | MA5-15738-D488 |
| Rabbit-anti-HA | No | ThermoFisher | PA1-985 |

**Table S2. Microscope settings for imaging channels.**

| Channel | λ excitation | Emission filter |
|---|---|---|
| roGFP blue | 405 nm | 525/50 |
| roGFP green | 445 nm | 525/50 |
| mGold | 515 nm | 542/27 |
| TRaQ-G | 561 nm | 642 LP |
| Alexa Fluor 488 | 488 nm | 525/50 |
| Alexa Fluor 680 | 561 nm | 642 LP |

**Table S3. Plasmid sources**

| Plasmid | Source | Vector | Insert |
|---|---|---|---|
| Calnexin-Halo-mGold | Addgene 183989 | - | - |
| pmTurquoise2-Golgi | Addgene 36205 | - | - |

| | | | |
|---|---|---|---|
| B4GALT1-Grx1-roGFP1-iE | TwistBioscience | - | - |
| The B4GALT1-Grx1-roGFP1-iE gene sequence is available in Addgene cloned into a pCMV backbone (to be added) | | | |
| HA-sCGrx1p-6His | TwistBioscience | - | - |
| sCGrx1pER | A gift from Appenzeller-Herzog[16] | - | - |
| B4GALT1-HA-sCGrx1p | Gibson assembly | B4GALT1-pmTurquoise2 | sCGrx1pER[16] |
| B4GALT1-sCGrx1p-HA | Gibson assembly (Addgene -to be added) | B4GALT1-HA-sCGrx1p | B4GALT1-HA-sCGrx1p and B4GALT1-HA-sCGrx1p |
| HA-B4GALT1-sCGrx1p | Gibson assembly | B4GALT1-HA-sCGrx1p | B4GALT1-HA-sCGrx1p B4GALT1-HA-sCGrx1p |
| HT7-mGold-6His | Addgene 183986 | | |
| B4GALT1-HT7-mGold | Gibson assembly (Addgene -to be added) | B4GALT1-pmTurquoise2 | Calnexin-Halo-mGold |

**Table S4. Primers for Gibson assembly**

| Plasmid | Fragments | Primers | |
|---|---|---|---|
| B4GALT1-HA-sCGrx1p | Vector: B4GALT1-pmTurquoise2 | For | TTGCAAATtaaagcggccgcgact |
| | | Rev | ATACGGATAcaccatggtggcgacc |
| | Insert: SCGRX1P-er | For | accatggtgTATCCGTATGATGTGCCTGACTACGC |
| | | Rev | gccgctttaATTTGCAAGAATAGGTTCTAACAATTCCTCC |
| HA-B4GALT1-sCGrx1p | Vector: B4GALT1-ha-scgrx1p | For | cgccaccGTATCTCAAGAAACTATCAAGCACGTCAAG |
| | | Rev | ATACGGATAcatggtggcgagct |
| | Insert 1: galt1-HA-scgrx1p | For | gccaccatgTATCCGTATGATGTGCCTGAC |
| | | Rev | ccgaagcctTTCTTGAGATACCATGGTACCTTCTGC |
| | Insert 2: galt1-ha-SCGRX1P | For | TCAAGAAaggcttcgggagccg |
| | | Rev | TGAGATACggtggcgaccggtgg |
| B4GALT1- | Vector: B4GALT1-ha-scgrx1p | For | ACGCAGAAGGTACCtaaagcggcc |
| | | Rev | GTACCTTCggtggcgaccggtggat |
| | Insert 1: | For | TGCAAATgtcgccaccatggtgTATCC |

| | | | |
|---|---|---|---|
| sCGrx1p-HA | galt1-HA-scgrx1p | Rev | gccgctttaGGTACCTTCTGCGTAGTCAGG |
| | Insert 2: galt1-ha-SCGRX1P | For | cgccaccGAAGGTACCATGGTATCTCAAGAAACTATCAAG |
| | | Rev | gtggcgacATTTGCAAGAATAGGTTCTAACAATTCCTCC |
| B4GALT1-HT7-mGold | pmTurquoise2-Golgi | For | aagtaaagcggccgc |
| | | Rev | atttcggtggcgacc |
| | Calnexin-Halo-mGold | For | cgccaccgaaatcggtact |
| | | Rev | cggccgctttacttgtacag |

**Table S5:** Important sequences of used GOIs for mammalian cell expression.

| Plasmid | GOI sequence |
|---|---|
| B4GALT1 | aggcttcgggagccgctcctgagcggcagcgccgcgatgccaggcgcgtccctacagcgggcctgccgcct<br>gctcgtggccgtctgcgctctgcaccttggcgtcaccctcgtttactacctggctggccgcgacctgagccgcctg<br>ccccaactggtcggagtctccacaccgctgcag |
| Grx1-roGFP2 | atggctcaagagtttgtgaactgcaaaatccagcctgggaaggtggttgtgttcatcaagcccacctgcccgtac<br>tgcaggagggcccaagagatcctcagtcaattgcccatcaaacaagggcttctggaatttgtcgatatcacagc<br>caccaaccacactaacgagattcaagattatttgcaacagctcacgggagcaagaacggtgcctcgagtcttt<br>attggtaaagattgtataggcggatgcagtgatctagtctctttgcaacagagtggggaactgctgacgcggcta<br>aagcagattggagctctgcagactagtggtggttcaggtggtggtggttcaggtggtggtggttcaggtggagga<br>ggatcaggaggaggaggatcaggaggaggaggatcaggaggagaattcgtgagcaagggcgaggagct<br>gttcaccggggtggtgcccatcctggtcgagctggacggcgacgtaaacggccacaagttcagcgtgtccggc<br>gagggcgagggcgatgccacctacggcaagctgaccctgaagttcatctccaccaccggcaagctgcccgt<br>gccctggcccaccctcgtgaccaccctgacctacggcgtgcagtgcttcagccgctaccccgaccacatgaag<br>cagcacgacttcttcaagtccgccatgcccgaaggctacgtccaggagcgcaccatcttcttcaaggacgacg<br>gcaactacaagacccgcgccgaggtgaagttcgagggcgacaccctggtgaaccgcatcgagctgaaggg<br>catcgacttcaaggaggacggcaacatcctggggcacaagctggagtacaactacaactgccacaacgtct<br>atatcatggccgacaagcagaagaacggcatcaaggtgaacttcaagatccgccacaacatcgaggacgg<br>cagcgtgcagctcgccgaccactaccagcagaacacccccatcggcgacggccccgtgctgctgcccgaca<br>accactacctgagcacctgctccgccctgagcaaagaccccaacgagaagcgcgatcacatggtcctgctgg<br>agttcgtgaccgccgccgggatcactctcggcatggacgagctgtacaag |
| Grx1-roGFP1-iE | atggctcaagagtttgtgaactgcaaaatccagcctgggaaggtggttgtgttcatcaagcccacctgcccgtac<br>tgcaggagggcccaagagatcctcagtcaattgcccatcaaacaagggcttctggaatttgtcgatatcacagc<br>caccaaccacactaacgagattcaagattatttgcaacagctcacgggagcaagaacggtgcctcgagtcttt<br>attggtaaagattgtataggcggatgcagtgatctagtctctttgcaacagagtggggaactgctgacgcggcta<br>aagcagattggagctctgcagactagtggtggttcaggtggtggtggctcgggtggtggtggatcaggaggag<br>gcggtagtggaggaggtggaagcggcggaggaggatcaggaggagaattcgtgagcaagggcgaggag<br>ctgttcaccggggtggtgcccatcctggtcgagctggacggcgacgtaaacggccacaagttcagcgtgtccg<br>gcgagggcgagggcgatgccacctacggcaagctgaccctgaagttcatctccaccaccggcaagctgccc<br>gtgccctggcccaccctcgtgaccaccctgagctacggcgtgcagtgcttcagccgctaccccgaccacatga<br>agcagcacgacttcttcaagtccgccatgcccgaaggctacgtccaggagcgcaccatcagcttcaaggacg<br>acggcaactacaagacccgcgccgaggtgaagttcgagggcgacaccctggtgaaccgcatcgagctgaa<br>gggcatcgacttcaaggaggacggcaacatcctggggcacaagctggagtacaactacaactgcgagagc<br>aacgtctatatcaccgccgacaagcagaagaacggcatcaaggctaacttcaagacccgccacaacatcga<br>ggacggcagcgtgcagctcgccgaccactaccagcagaacacccccatcggcgacggccccgtgctgctg<br>cccgacaaccactacctgagcacctgctccgccctgagcaaagaccccaacgagaagcgcgatcacatggt<br>cctgctggagttcgtgaccgccgccgggatcactctcggcatggacgagctgtacataagctaa |
| sCGrx1p-HA | atggtatctcaagaaactatcaagcacgtcaaggaccttattgcagaaaacgagatcttcgtcgcatccaaaac<br>gtactgtccatactctcatgcagctttaaacacgctttttgaaaagttaaaggttcccaggtccaaagttctggttttg<br>caattgaatgacatgaaggaaggcgcagacattcaggctgcgttatatgagattaatggccaaagaaccgtgc<br>caaacatctatattaatggtaaacatattggaggcaacgacgacttgcaggaattgagggagactggtgaattg<br>gaggaattgttagaacctattcttgcaaatgtcgccaccatggtgtatccgtatgatgtgcctgactacgca |

| | |
|---|---|
| HT-mGold | gaaatcggtactggctttccattcgacccccattatgtggaagtcctgggcgagcgcatgcactacgtcgatgttg<br>gtccgcgcgatggcacccctgtgctgttcctgcacggtaacccgacctcctcctacgtgtggcgcaacatcatcc<br>cgcatgttgcaccgacccatcgctgcattgctccagacctgatcggtatgggcaaatccgacaaaccagacct<br>gggttatttcttcgacgaccacgtccgcttcatggatgccttcatcgaagccctgggtctggaagaggtcgtcctg<br>gtcattcacgactggggctccgctctgggtttccactgggccaagcgcaatccagagcgcgtcaaaggtattgc<br>atttatggagttcatccgccctatcccgacctgggacgaatggccagaatttgcccgcgagaccttccaggcctt<br>ccgcaccaccgacgtcggccgcaagctgatcatcgatcagaacgtttttatcgagggtacgctgccgatgggtg<br>tcgtccgcccgctgactgaagtcgagatggaccattaccgcgagccgttcctgaatcctgttgaccgcgagcca<br>ctgtggcgcttcccaaacgagctgccaatcgccggtgagccagcgaacatcgtcgcgctggtcgaagaatac<br>atggactggctgcaccagtcccctgtcccgaagctgctgttctggggcaccccaggcgttctgatcccaccggc<br>cgaagccgctcgcctggccaaaagcctgcctaactgcaaggctgtggacatcggcccgggtctgaatctgctg<br>caagaagacaacccggacctgatcggcagcgagatcgcgcgctggctgtccacgctcgagatttccggcgtg<br>agcaagggcgaggagctgttcaccggggtggtgcccatcctggtcgagctggacggcgacgtaaacggcca<br>caagttcagcgtgtccggcgagggcgagggcgatgccacctacggcaagctgaccctgaagttcatctgcac<br>caccggcaagctgcccgtgccctggcccaccctcgtgaccagcctgggctacggcctgcagtgcttcgcccgc<br>taccccgaccacatgaagcagcacgacttcttcaagtccgccatgcccgaaggctacgtccaggagcgcacc<br>atcttcttcaaggacgacggcaactacaagacccgcgccgaggtgaagttcgagggcgacaccctggtgaa<br>ccgcatcgagctgaagggcatcgacttcaaggaggacggcaacatcctggggcacaagctggagtacaact<br>acaacagccacaacgtctatatcaccgccgacaagcagaagaacggcatcaaggccaacttcaagatccg<br>ccacaacatcgaggacggcggcgtgcagctcgccgaccactaccagcagaacacccccatcggcgacgg<br>ccccgtgctgctgcccgacaaccactacctgagctaccagtccaagctgagcaaagaccccaacgagaagc<br>gcgatcacatggtcctgctggagttcgtgaccgccgccgggatcactctcggcatggacgagctgtacaag |

### 2. Supplementary figures

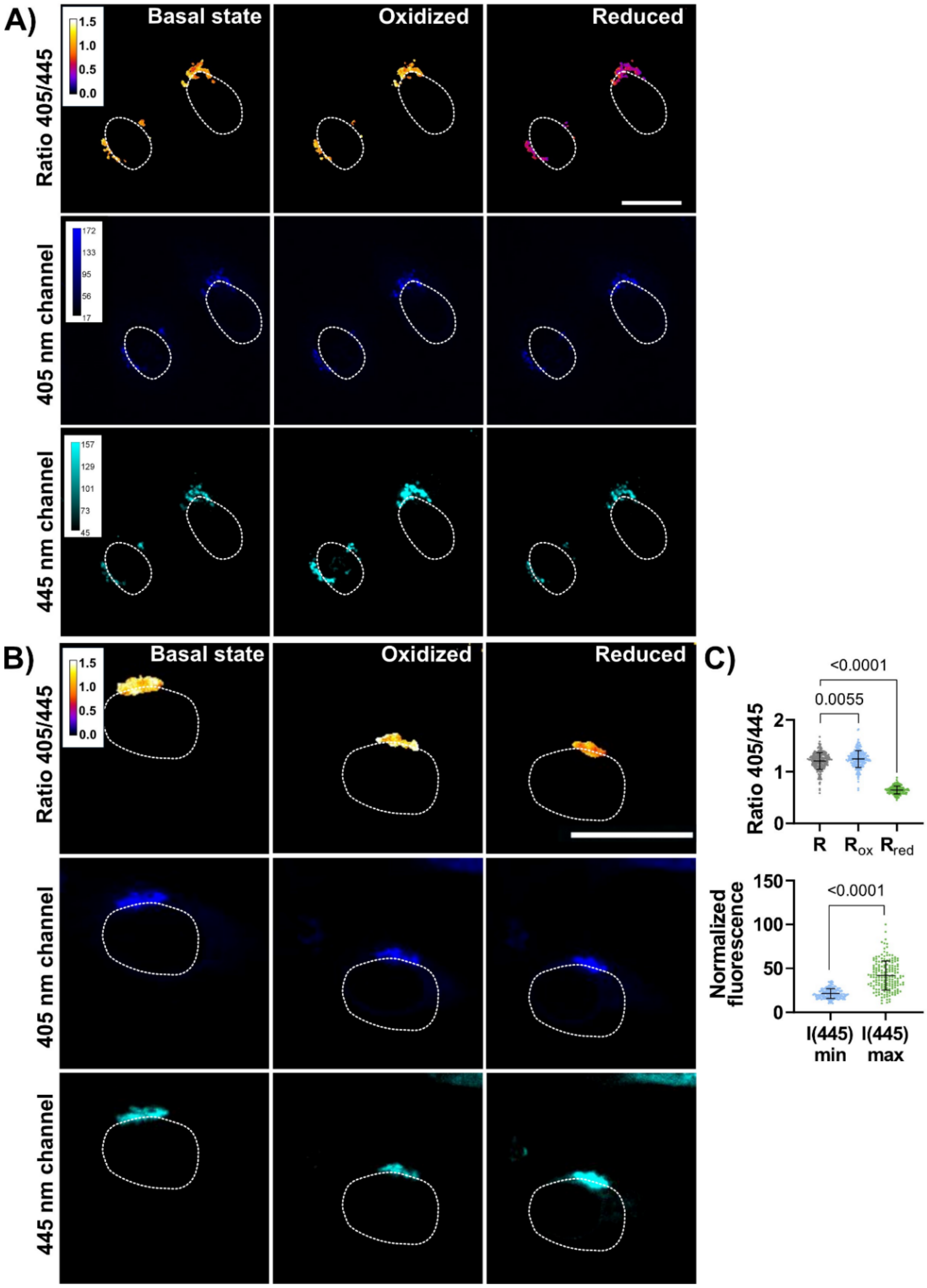


**Figure S1.** HeLa **(A)** and HEK293 (**B** and **C**) cells transiently transfected with B4GALT1-Grx1-roGFP1-iE. **A** and **B)** 405 nm channel, 445 nm channel, and ratiometric fluorescent images with excitation at 405 nm and 445 nm wavelength and emission at 525/50 nm. The cells were imaged at basal state conditions, under oxidizing conditions (after 3 min of adding 1 mM $H_2O_2$), and under reducing conditions (after 3 min of adding 5 mM DTT). Scale bar = 20 µm. **C)** Data analysis of n ≥ 50, technical replicates, average of three biological replicates. Statistical significance was assessed by unpaired, two-tailed, Mann-Whitney test.

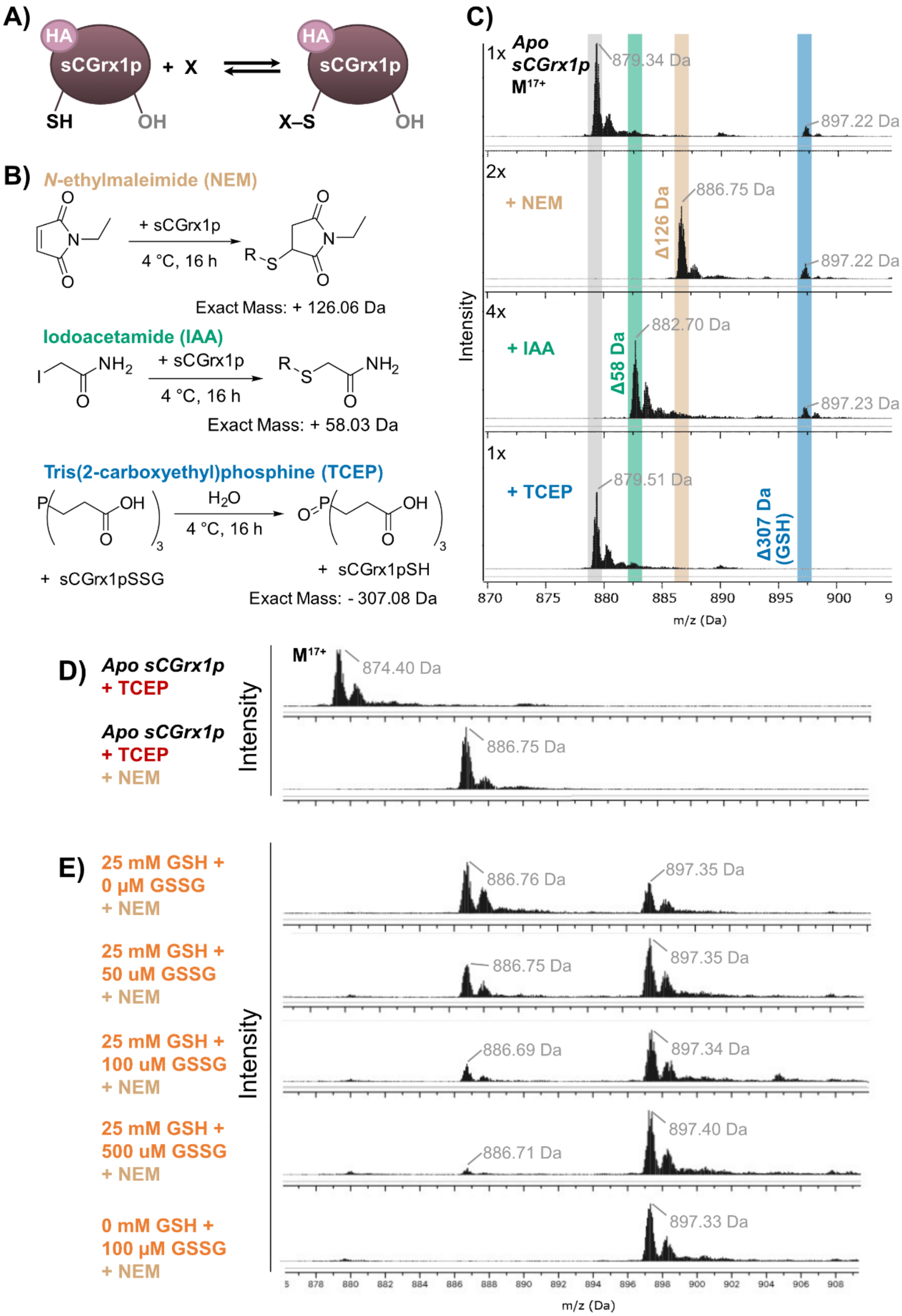


**Figure S2.** sCGrx1p *in vitro* alkylation assays analyzed via protein MS. **A)** Schematic reaction of cysteine alkylation. **B)** Reaction scheme of sCGrx1p alkylation with NEM and IAA, and reduction with TCEP. **C)** HR-ESI-UPLC-MS zoomed-in spectra of the different samples with indications of the expected mass changes upon chemical modification. All the reactions were performed in PBS (100 µM protein and 2 mM reagent for 4 °C, 16 h). **D)** HR-ESI-MS zoomed in spectra of the apo-protein samples (100 µM) with prereduction with TCEP (0.5 mM, storage buffer) and NEM reaction (50 mM, 4 °C, 16 h). **E)** HR-ESI-MS zoomed-in spectra of protein samples incubated with GSH and increasing amounts of GSSG. The protein (100 µM) was pre-reduced with TCEP (0.5 mM, storage buffer), and the buffer was exchanged to PBS pH 7.2. The reduced protein was reacted with GSH/GSSG (20 °C, 10 min) and immediately capped by reaction with NEM (50 mM, 4 °C, 16 h).

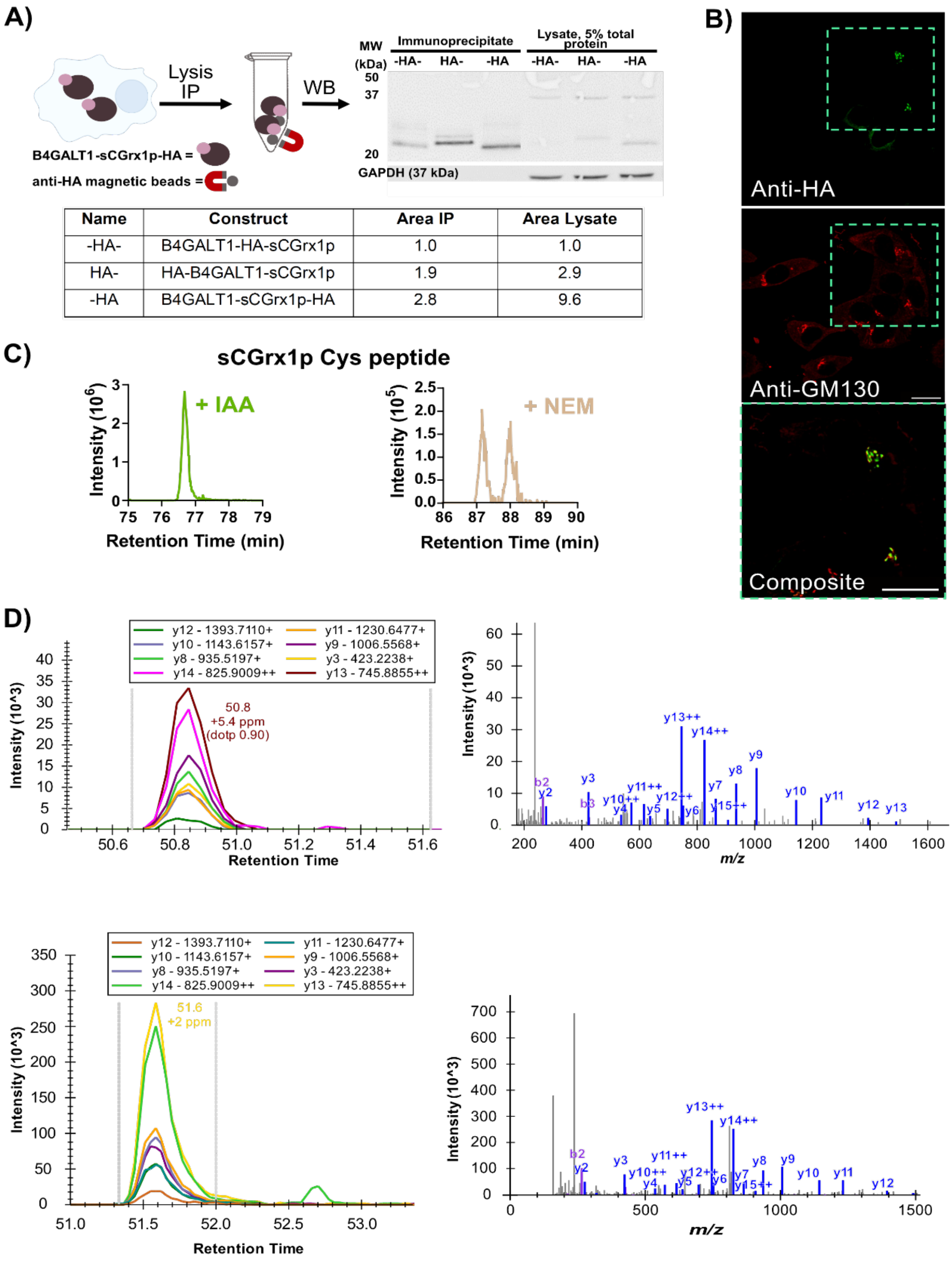



| Name | Construct | Area IP | Area Lysate |
|---|---|---|---|
| -HA- | B4GALT1-HA-sCGrx1p | 1.0 | 1.0 |
| HA- | HA-B4GALT1-sCGrx1p | 1.9 | 2.9 |
| -HA | B4GALT1-sCGrx1p-HA | 2.8 | 9.6 |

**Figure S3. A)** Schematic IP procedure and western blot analysis of the HA-tagged constructs transiently expressed in HeLa cells. Shown are the immunoprecipitates (after incubation at 4 °C for 16 h) from anti-HA magnetic beads (left) and raw lysate samples containing 5% of total protein (right). Bottom, Quantification of HA availability for HA IP. GAPDH was used as a control for overall expression levels. Areas were measured with Fiji and normalized regarding the -HA- construct. An additional normalization was carried out to the expression of GAPDH. **B)** Fluorescence microscopy images of fixed HeLa cells transiently expressing B4GALT1-sCGrx1p-HA. Mouse anti-HA and anti-mouse-Alexa488 antibodies were used to label the construct, and rabbit anti-GM130 and anti-rabbit-Alexa680 were used to label the Golgi marker

GM130. **C)** Extracted ion chromatogram (XIC) traces in MS1 of the sCGrx1p peptide containing the reactive cysteine and the alkylating modification (TY<u>C</u>PYSHAALNTLFEK, 638.9733 Da [M]+++ for IAA, 661.6487 Da [M]+++ for NEM). **D)** Fragment ion traces in Skyline for HeLa (top) and HEK293 (bottom) and MS/MS spectra of the peptide containing the modification. Y14++ was chosen as the quantification fragment since it is the most abundant fragment containing the IAA modification.

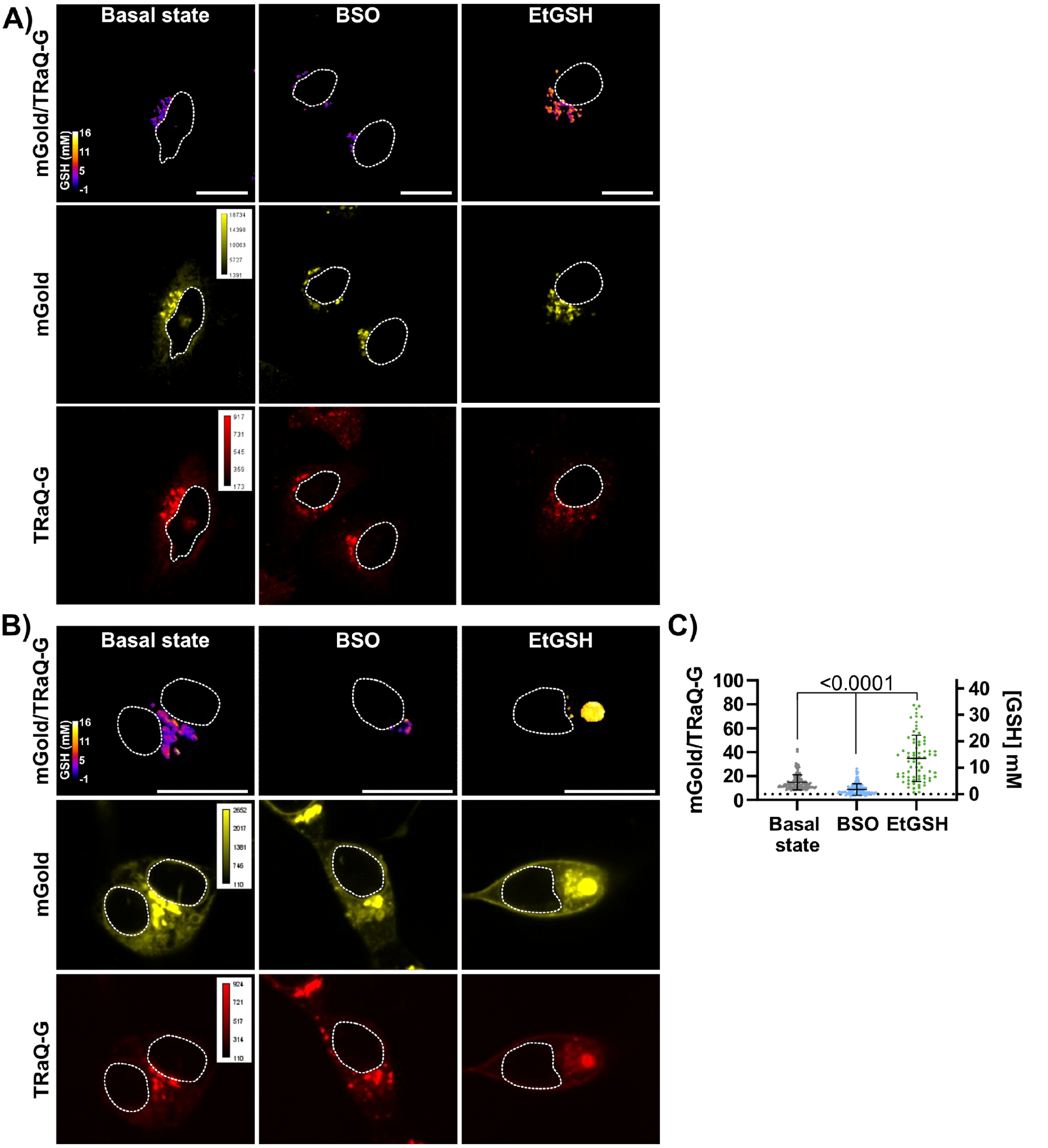


**Figure S4.** HeLa (A) and HEK293 (B and C) cells transiently transfected with B4GALT1-HT-mGold and incubated with 100 nM TRaQ-G ligand for 1 h at 37 °C. **A)** and **B)** mGold channel, TRaQ-G channel, and mGold/TRaQ-G ratiometric images of HeLa (A) and HEK293 (B) cells. Excitation at 515 nm and emission at 542/27 nm for mGold, and excitation at 561 nm and emission at 642 long-pass filter for TRaQ-G. The color calibration is illustrated in concentration values, not ratios. The cells were imaged at basal state conditions, under BSO treatment (1 mM, 3–4h incubation) and under EtGSH conditions (10 mM, 3–4h incubation). Scale bar = 20 µm. **C)** Data analysis of $n \geq 50$, technical replicates (individual cells), average of three biological replicates. Statistical significance was assessed by unpaired, two-tailed, Mann-Whitney test.
